# Correcting for Background Noise Improves Phenotype Prediction from Human Gut Microbiome Data

**DOI:** 10.1101/2021.03.19.436199

**Authors:** Leah Briscoe, Brunilda Balliu, Sriram Sankararaman, Eran Halperin, Nandita R. Garud

## Abstract

The ability to predict human phenotypes accurately from metagenomic data is crucial for developing biomarkers and therapeutics for diseases. However, metagenomic data is commonly affected by technical or biological variables, unrelated to the phenotype of interest, such as sequencing protocol or host sex, which can greatly reduce or, when correlated to the phenotype of interest, inflate prediction accuracy. We perform a comparative analysis of the ability of different data transformations and existing supervised and unsupervised methods to correct microbiome data for background noise. We find that supervised methods are limited because they cannot account for unmeasured sources of variation. In addition, we observe that unsupervised approaches are often superior in addressing these issues, but existing methods developed for other ‘omic data types, e.g., gene expression and methylation, are restricted by parametric assumptions unsuitable for microbiome data, which is typically compositional, highly skewed, and sparse. We show that application of the centered log-ratio transformation prior to correction with unsupervised approaches improves prediction accuracy for many phenotypes while simultaneously reducing variance due to unwanted sources of variation. As new and larger metagenomic datasets become increasingly available, background noise correction will become essential for generating reproducible microbiome analyses.

## Introduction

The human gut microbiome is associated with a number of host phenotypes including colorectal cancer^1^, obesity^2,3^, and antibiotic consumption^4–7^, among other traits^8,9^. Despite the promise of the microbiome as a diagnostic tool, significant challenges remain for accurate prediction of human phenotypes from microbiome data. One major challenge is that several covariates are known to introduce unwanted variation and systematically bias the relative abundances of taxonomic units in microbiome samples^10–16^. These covariates include host diet and sex^17–19^, sample storage^20^, cell lysis protocol^10,21^, DNA preservation and storage protocol^22^, preparation kit^23^, extraction method^24,25^, and primer choice^21^. This variation can substantially reduce the ability to predict host phenotypes from microbiome data. In addition, when these variables are also correlated with the phenotype of interest, they can act as confounders, inducing associations between the microbiome and the phenotype and artificially inflating phenotypic prediction accuracy. For example, confounding covariates were found to be pervasive^26^ in one of the largest metagenomic datasets available, the American Gut Project (AGP)^27^. Confounding can also arise when datasets from different studies are combined together to augment the power to detect associations^8,28^, a practice that is becoming increasingly common^29–33^ and is potentially a powerful means to validate results in a discovery dataset with held out datasets^1,34,35^. For example, Gibbons *et al*.^36^ found that combining datasets to detect members of the microbiome that were associated with colorectal cancer resulted in false positive detection of differentially abundant taxa due to the different case-control ratio in different combined studies.

Despite the widespread effects of background noise in microbiome data, there is currently a dearth of methods specially equipped for removing unwanted variation in microbiome data. Existing methods repurposed from other domains, including gene expression^39,40^ and methylation^41,42,37–39^, generally fall into two categories: supervised methods, where the sources of variation must be explicitly specified, and unsupervised methods, where the sources of variation are first inferred and then removed before association or prediction. The most popular supervised methods are batch mean centering (BMC)^43^, which centers data batch by batch, and ComBat^44^ and limma^45^, which both use empirical Bayes. However, since many sources of variation may be unknown, and moreover, the extent of variation they introduce may vary from dataset to dataset^13,36,40–42^, unsupervised approaches^43–45^ for covariate correction may be more effective in removing background noise.

Despite their promise, the repurposed supervised and unsupervised approaches^43–45^ are not suitable for microbiome data because they rely on assumptions that the data is normally distributed. However, taxonomic features are often sparse^46,47^ due to taxa having abundances below the detection limit of sequencing^46^, or taxa being absent in certain samples, resulting in skewed non-normal distributions. Additionally, microbiome data is compositional, or, in other words, typically represented as a relative abundance table in which the frequencies of taxonomic units within a sample sum to one. This induces correlations between taxonomic units within a sample. Recently, supervised methods were proposed explicitly for microbiome data, and include percentile normalization^31^, Partial Least Squares Discriminant Analysis^48^, and multiplicative bias correction^15^. Both percentile normalization^31^ and Partial Least Squares Discriminant Analysis^48^ aim to find predictive features in fully labeled data with known batches and known phenotypes, and are not designed for prediction of phenotypes in unlabeled data, while multiplicative bias correction^15^ requires a reference sample in which the species abundance distribution is known, and thus cannot be applied to the majority of microbiome data that lack such references. Given that these methods are supervised and thus cannot be applied to unlabeled data, there still remains a need in the microbiome field for unsupervised approaches that can adjust for both measured and unmeasured variables.

In this paper, we evaluated different supervised background noise correction approaches^49–51^ that are commonly used for microbiome data and compared them to an unsupervised approach in which the top principal components (PCs) from microbiome data are regressed from OTUs or *k*-mers (a reference-agnostic feature alternative to OTUs, based on short substrings of length *k* in raw sequences) after application of the centered log ratio (CLR). This PC correction approach has been used in other applications, such as correcting for population structure in genetic data^70–72^ and morphological differences in a study population of sticklebacks^73^, but to date has not been applied to microbiome data. CLR is widely used for compositional data and specifically in microbiome contexts^28,52–57^, and is a suggested transformation prior to factor analysis, such as principal components analysis (PCA), because it breaks the dependence between features while also ensuring that the resulting covariance matrix is non-singular^53^. CLR additionally makes data more normally distributed^48^. In this paper, we show that CLR transformation is a crucial step in revealing sources of background noise for proper covariate correction and phenotype prediction. We found that the combined approach of PC correction in conjunction with CLR can simultaneously address the compositionality and non-normality, while use of *k-*mers addresses sparsity, of microbiome data all while improving the ability to predict human phenotypes after correcting for background noise.

Throughout this study, we highlight important considerations for phenotype prediction from large cohort and cross-study analyses, which we hope paves the way for higher reproducibility across microbiome studies.

## Results

We analyzed three metagenomic datasets for evidence of biological and technical covariates that could introduce noise or confounding that, as a result, could diminish or inflate phenotype prediction accuracy. We evaluated the ability of three popular supervised approaches for microbiome data^49–51^ and an unsupervised approach, PCA correction, to correct for noise and confounding in these three datasets, and the impact of this correction on phenotype prediction accuracy. We focused on three phenotypes of interest: body mass index (BMI), antibiotic consumption, and colorectal cancer (CRC) (**Table 1**). The datasets we analyzed included: (i) the American Gut Project^27^ (AGP), which has known confounding variables^26^ (ii) a pooled dataset composed of three 16S datasets with healthy and CRC individuals (hereafter referred to as ‘CRC-16S’)^31^, and (iii) a pooled dataset composed of seven whole genome sequenced datasets (WGS) with healthy and CRC individuals (hereafter referred to as ‘CRC-WGS’)^1^. These datasets allowed us to assess noise and confounding both within a dataset (AGP) and across pooled datasets (CRC-16S and CRC-WGS).

**Table 1.** Datasets used in this study. Two pooled datasets composed of multiple studies are abbreviated as CRC-16S^58–60^ and CRC-WGS ^1,59,61–64^, whereas the American Gut Project (AGP)^27^ is composed from one source study but has several known confounders^26^.

| Phenotype | Joined dataset | Number of samples | Number of studies | Sequencing method | Published Sources |
| --- | --- | --- | --- | --- | --- |
| Body mass index | American Gut Project (AGP) | 16,225 | 1 ( multiple sequencing batches) | 16S | McDonald et al. |
| Antibiotic history | American Gut Project (AGP) | 21,862 | 1 ( multiple sequencing batches) | 16S | McDonald et al. |
| Colorectal Cancer | CRC-16S | 574 | 3 | 16S | Baxter et al.<br>Zeller et al.<br>Zackular et al. |
| Colorectal Cancer | CRC-WGS | 813 | 7 | WGS | Feng et al.<br>Yu et al.<br>Vogtman et al.<br>Hannigan et al.<br>Thomas et al.<br>Zeller et al. |

### Background noise due to technical and biological variables is widespread in microbiome data

While it is known that there are biases and confounders in microbiome data^26^, it is not well understood how these factors impact our ability to predict host phenotypes from metagenomic data. To assess the impact of biological and technical contributors on microbiome variability and microbiome-phenotype relationship, we performed PCA on CLR-transformed (see Methods) OTUs and *k-*mers derived from the raw metagenomic reads. We observed that the top two PCs separate out dataset effects and not the primary phenotype of interest, as shown in the CRC-WGS pooled dataset (**Figure 1**) and the other two datasets (**Figure S1**), suggesting that the phenotype signal is weak in the presence of heterogeneity.

**Figure 1.**
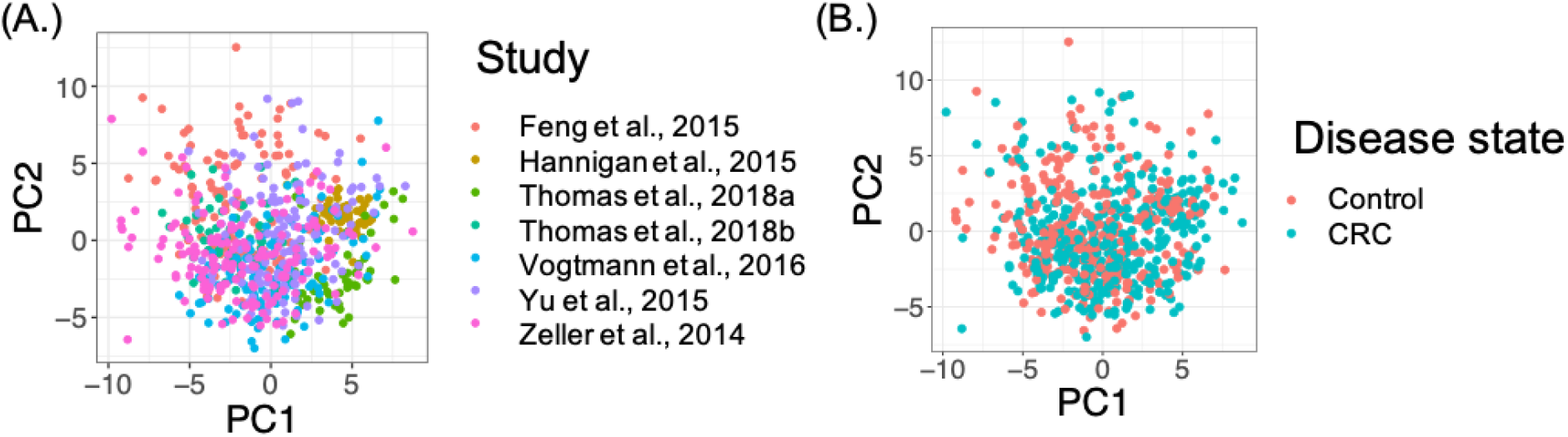
First two principal components from the CRC-WGS study. PCA was applied to OTU data from the CRC-WGS dataset, and samples were plotted along the first 2 PCs with colors in (A) indicating dataset membership and colors in (B) representing colorectal cancer status.

More generally, the top 10 PCs from OTUs (**Figure 2A and Figure S2**) were highly correlated with technical variables, e.g., dataset label has a Pearson correlation of 0.87 with PC2 in CRC-16S and 0.45 with PC2 in CRC-WGS, and, to a lesser extent, biological variables unrelated to the phenotype of interest, e.g., sex (CRC-WGS: *ρ*= 0.10 with PC5; CRC-16S: *ρ*= 0.11 with PC5) and age (AGP: *ρ*= 0.16 with PC2; CRC-WGS: *ρ* = 0.14 with PC2; CRC-16S: *ρ*= 0.16 with PC1). Other noteworthy correlations among the top 10 PCs include country of the host (CRC-WGS: *ρ*= 0.47 with PC2), DNA extraction kit (CRC-WGS: *ρ*= 0.42 with PC2), collection year (AGP: *ρ*= 0.48 with PC5), and sequencing instrument (AGP: *ρ*= 0.45 with PC5; CRC-16S: *ρ*= 0.56 with PC1). Unlike the many technical and biological covariates, the phenotypes of interests were poorly or moderately correlated with the top 10 PCs and include antibiotic consumption in the AGP (*ρ* = 0.12 with PC3) and CRC status (CRC-WGS: *ρ* =0.24 with PC3; CRC-16S: *ρ*=0.20 with PC4), and BMI in the AGP (*ρ* = 0.52 with PC5).

**Figure 2.**
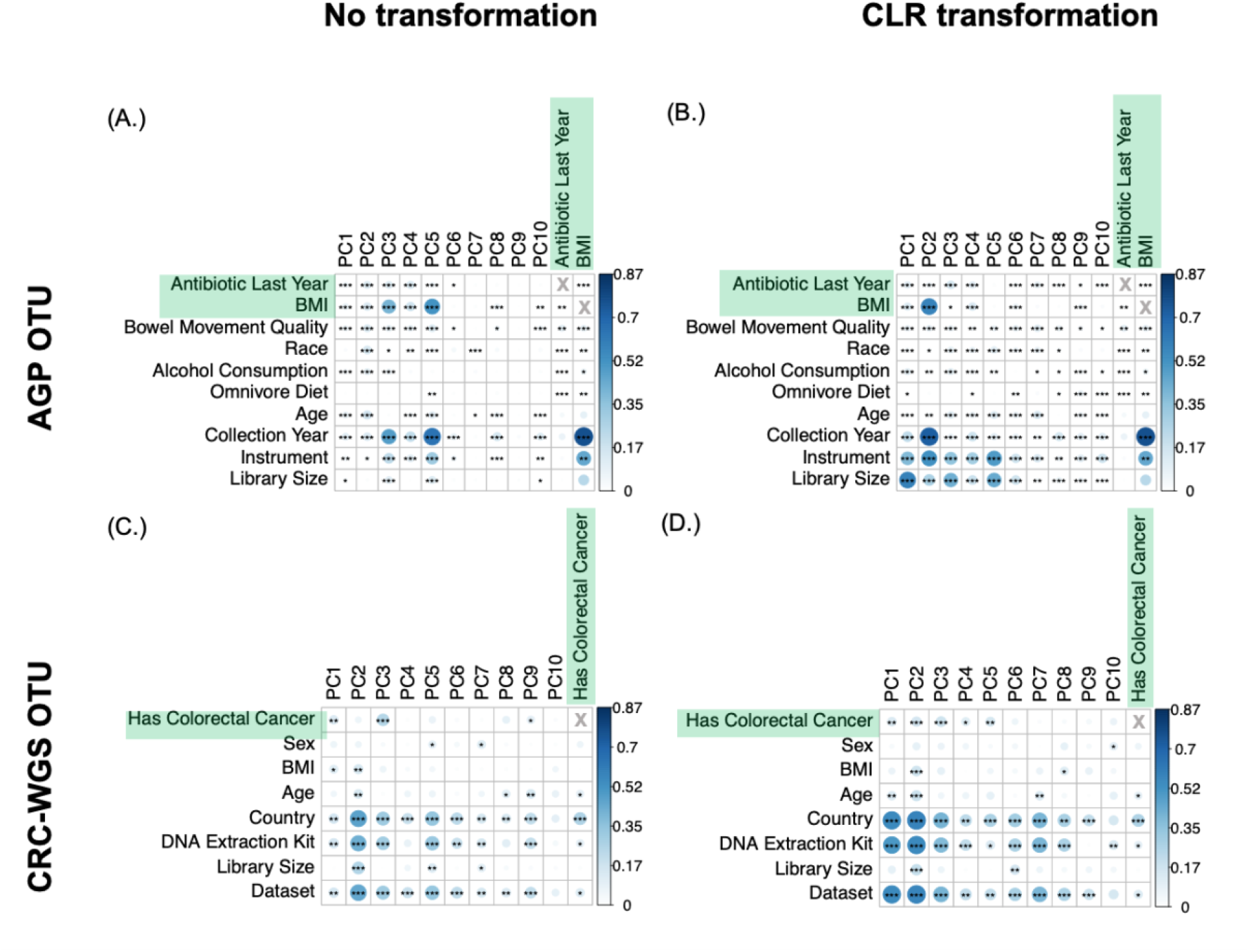
Microbiome data is affected by technical and biological variables. The first 10 PCs are correlated with variables measured in each of the studies, including dataset label, library size, DNA extraction kit used, country of origin, age, body mass index (BMI), sex, and colorectal cancer status (CRC). CLR reveals stronger correlations between the top principal components and confounding covariates compared to no transformation. The size and color of the circles in each cell indicates the magnitude of correlation and black asterisks indicate the significance of the Pearson correlation of the PCs with each of the variables. The color bar at right of each plot represents the range of correlations observed across all datasets. [*,**,*** indicate *p*-values as follows: 10^−2^ < *p* < 0.05, 10^−3^ < *p* < 10^−2^, 10^−4^ < *p* < 10^−3^, *p* < 10^−4^]. See **Figures S2** and **S5** for similar analyses for the CRC-16S joined dataset and 7-mers.

In addition to contributing a large proportion of microbiome variance, the phenotypes of interest were significantly correlated with technical and biological covariates, which means that these covariates could potentially confound the ability to predict phenotypes. For example, the country of host, DNA extraction kit, and dataset were significantly correlated to colorectal cancer status in CRC-WGS (*ρ* =0.26; *ρ* = 0.10, and *ρ* = 0.13, respectively) and sequencing instrument and collection year in the AGP were significantly correlated with BMI (*ρ* =0.45 and *ρ* = 0.76, respectively).

Compared to no transformation, CLR transformation resulted in more normally distributed data (**Figure S3**), and stronger correlations between each of the top 10 PCs and phenotypes of interest and technical covariates (**Figure 2B, 2D** and **S2**). For example, without CLR transformation, antibiotic consumption in AGP was significantly correlated with the first 6 PCs that collectively explain 33% of microbiome variance (p-value ≤ 0.02 for each PC significantly correlated, where the first 10 PCs were tested for correlations). However after CLR, antibiotic consumption was significantly correlated with all the first 10 PCs that collectively explain 50% of microbiome variance (p ≤ 0.01 for each of the 10 PCs tested). Similarly, CRC status in CRC-WGS was initially significantly correlated with PCs 1, 3 and 9 (p ≤ 0.03 for each PC) that collectively explain 17% of microbiome variance, but after CLR, CRC status was significantly correlated with PCs 1 through 5 (p ≤ 0.01 for each PC) that collectively explain 31% of microbiome variance. Similar trends were found with BMI in AGP (**Figure 2B**) but not with CRC in CRC-16S **(Figure S2**), although correlations significantly increase in magnitude for all three datasets across all covariates **Figure S4**). Technical variables were more correlated with top PCs than phenotypes of interest after CLR: the strongest correlations in CLR-transformed AGP, CRC-WGS, and CRC-16S pooled datasets were between the first two PCs (which explain 14%, 12%, 15% of microbiome variance, respectively) and the dataset label (CRC-WGS: *ρ*= 0.57 with PC2; CRC-16S: *ρ*= 0.87 with PC1) and sequencing protocol (AGP: *ρ* = 0.52 with PC2; CRC-16S: *ρ*= 0.59 with PC1).

We also assessed the impact of *k*-merization on the correlation of variables with top PCs. Similar to OTUs, CLR transformation of *k*-mers results in more significant correlations compared to no transformation. In fact, the correlations of technical variables with PCs were stronger for *k*-mers than OTUs for all three datasets (**Figure S5**). For example, sequencing method is significantly correlated with CRC-16S PCs that collectively explained 21% of variance in OTUs, but 49% of variance in *k*-mers. PCs significantly correlated with CRC status in CRC-16S, by contrast, only collectively explained 16 and 18% of variance in OTUs and *k*-mers, respectively.

### Background noise correction increases variance explained by phenotype while reducing variance explained by confounders

We assessed how successfully background noise correction methods can remove variation due to technical and biological confounders without sacrificing the phenotype of interest, as this represents the net benefit of applying background noise correction (**Figure 3A** and **3B, Figure S6**). We focused on *k*-mers for this analysis because they show the most sensitivity to technical confounders. The supervised background noise correction methods we applied were BMC^49^, ComBat^51^, limma^50^. When testing BMC^49^, ComBat^51^ and limma^50^, we correct for the primary contributor of heterogeneity in each dataset: sequencing instrument and source study in the AGP and CRC datasets respectively, since these variables were the most correlated with the top PCs (**Figure 2**).

**Figure 3.**
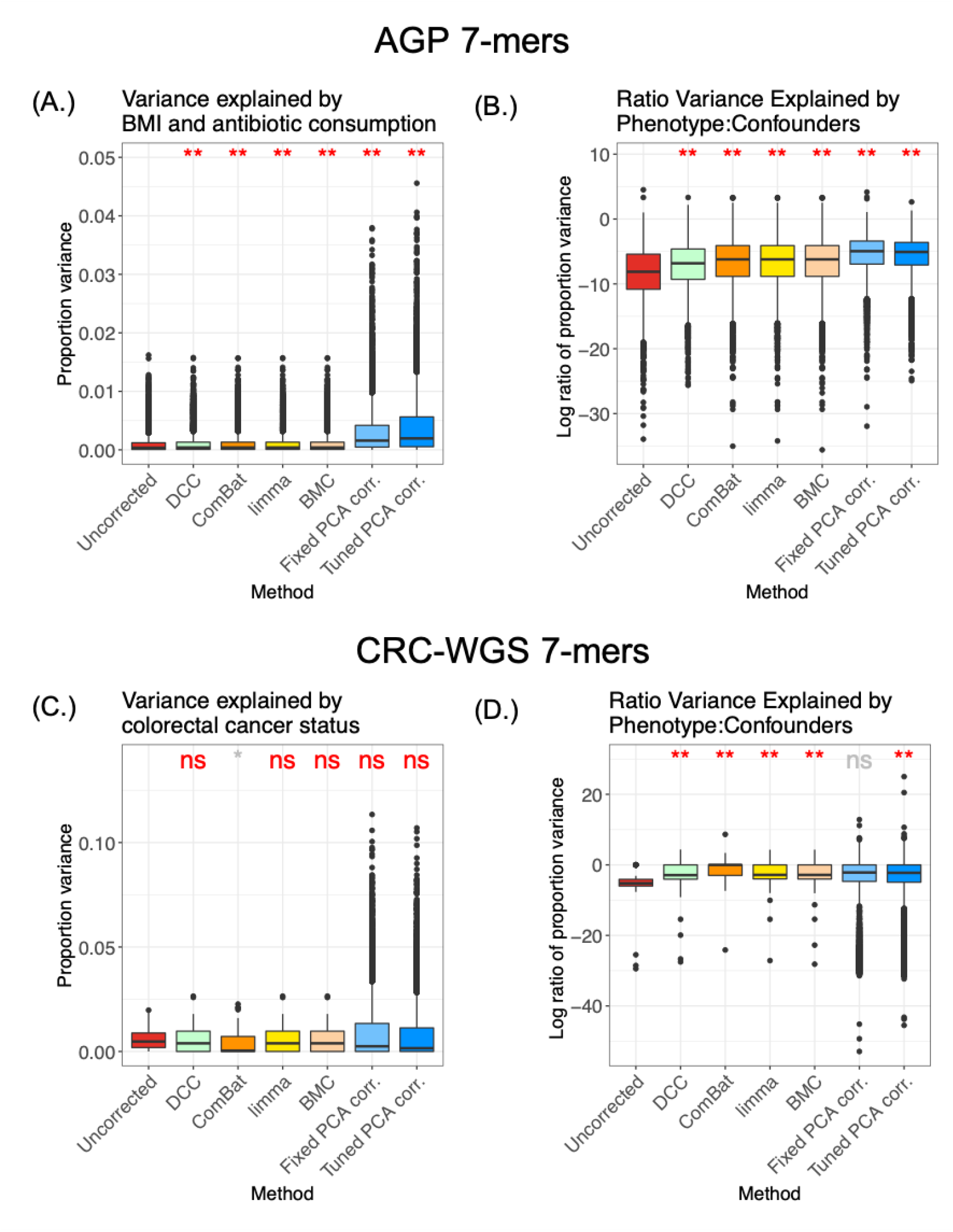
Proportion of variance explained by phenotypes of interest versus confounders. We plotted the distribution of the proportion of variance in each of the 7-mer features that is explained by (a) antibiotic consumption and BMI in the AGP cohort, and by (c) colorectal cancer status in the CRC-WGS joined datasets. We plotted the log2 ratio of that same proportion by the proportion explained by confounders, including DNA extraction kit, sequencing instrument, and sex in the (b) AGP cohort and (d) the CRC-WGS joined datasets. Each point represents one feature in the 7-mer matrix table. [ns,*,**,*** indicate *p*-values as follows: non-significant at 0.05 threshold, 10^−2^ < *p* < 0.05, 10^−3^ < *p* < 10^−2^, 10^−4^ < *p* < 10^−3^, *p* < 10^−4^, respectively, in a Wilcoxon rank sum test comparing each method to the baseline]. All methods’ baseline was Uncorrected except fixed-PCA, which was compared to DCC as a baseline. Refer to **Figure S6** for the equivalent analysis for WGS-16S.

Additionally, we assessed the ability of two variants of unsupervised correction approaches in which PCA correction is applied after CLR: one in which data is corrected for the number of PCs that maximizes phenotype prediction accuracy and another in which data is corrected for a fixed and arbitrary number of PCs. We refer to these two variants as tuned-PCA and fixed-PCA, respectively (see Methods). Tuned-PCA uses a validation set to determine the optimal number of PCs that maximize prediction accuracy while fixed-PCA correction corrects for the first three PCs. The choice of three PCs for this analysis was arbitrarily selected to avoid completely throwing away the signal associated with the phenotype of interest. We analyzed the effect of higher numbers of PCs in the supplement (**Figure S7**). We included the fixed-PCA approach in order to evaluate the ability of a correction method agnostic to the phenotype signal. Finally, we also applied a supervised correction in which the primary contributors of heterogeneity were regressed out, an approach we term in this paper as Direct Covariate Correction (DCC) (see Methods). Because it is the only method that explicitly adjusts for the primary potential confounders, DCC serves as a baseline assessing fixed-PCA correction, which may not fully correct these primary confounders but on the other hand may correct for additional confounders.

The proportion of variance explained by phenotype after applying each method is quantified by modeling the *k*-mers as an outcome in a linear mixed model that jointly considers all measured variables both biological and technical, and then estimating the fraction of variance explained by each of these variables (see Methods). Tuned and fixed PCA correction resulted in the greatest increase in proportion of variance explained by phenotypes of interest in all datasets relative to no correction or DCC, respectively, and also relative to all other methods (**Figure 3** and **Figure S6**). By contrast, limma^50^ and BMC^49^, produced a minimal, albeit significant, increase in the proportion of variance explained by the phenotypes of interest in AGP and insignificant increase or decrease in variance explained by CRC in CRC-WGS and CRC-16S. In the case of CRC-WGS and CRC-16S, ComBat^51^ significantly reduces the variance that can be explained by CRC compared to uncorrected data (p < 0.05) (**Figure 3C** and **Figure S6**).

Additionally, we computed the ratio of variance explained by the phenotype of interest versus variance explained by the confounders (i.e., sequencing instrument, collection year, race, diet, alcohol consumption, type of bowel movement, library size, and age). Compared to uncorrected data, the tuned-PCA correction approach resulted in an increase in mean ratio by a factor of 6.9 and 1.2 (**Figures 3B, 3D, and S6**) for the AGP and CRC-WGS datasets, respectively, but resulted in a reduction for CRC-16S by a factor of 0.74. Fixed-PCA performed better than DCC in AGP by a factor of 3.5, but performed worse than DCC in CRC-WGS and CRC-16S by factors of 0.49 and 0.44, respectively (**Figures 3B, 3D, and S6**). In AGP and CRC-WGS, ComBat^51^, limma^50^, and BMC^49^ also had significantly increased ratios, but tuned PCA performed the best. In CRC-16S, only ComBat^51^ resulted in an increased ratio compared to uncorrected data (increased by a factor of 1.3) (**Figure S6**).

We conclude from these findings that existing background noise correction methods addressed variance due to confounders at the cost of the signal relevant to the phenotype of interest. Both the fixed and tuned-PCA correction were able to increase the variance attributed to phenotypes of interest across all datasets for the majority of features.

### Impact of data transformations and background noise correction on phenotype prediction

We evaluated the impact of each correction approach on prediction of a host’s colorectal cancer status, antibiotic consumption, and BMI when using either OTUs or *k*-mers as input features to a Random Forest classifier or linear regression model (see Methods). We hypothesized that, in scenarios when phenotype is not directly correlated to other covariates, background noise correction approaches will yield better prediction than uncorrected data. In cases where covariates confound the phenotype directly, these approaches may not improve, and even reduce, phenotype prediction accuracy because correction for confounders would also result in removal of the phenotype of interest.

Among OTUs, all methods^49–51^ either had no effect or significantly reduced accuracy for all phenotypes, compared to no correction (**Figure 4**). The reduction in accuracy is expected when phenotypes of interest are confounded by covariates, as observed in previous sections. Notably, fixed-PCA correction resulted in the greatest decrease in phenotype prediction accuracy, even relative to its comparable baseline, DCC. **Figure S7** shows that the AUC for phenotype prediction increases monotonically as the first three PCs are regressed out. However, if too many PCs are regressed out, then, phenotype prediction accuracy starts to decrease indicating PCs associated with the phenotypic signal were removed.

**Figure 4.**
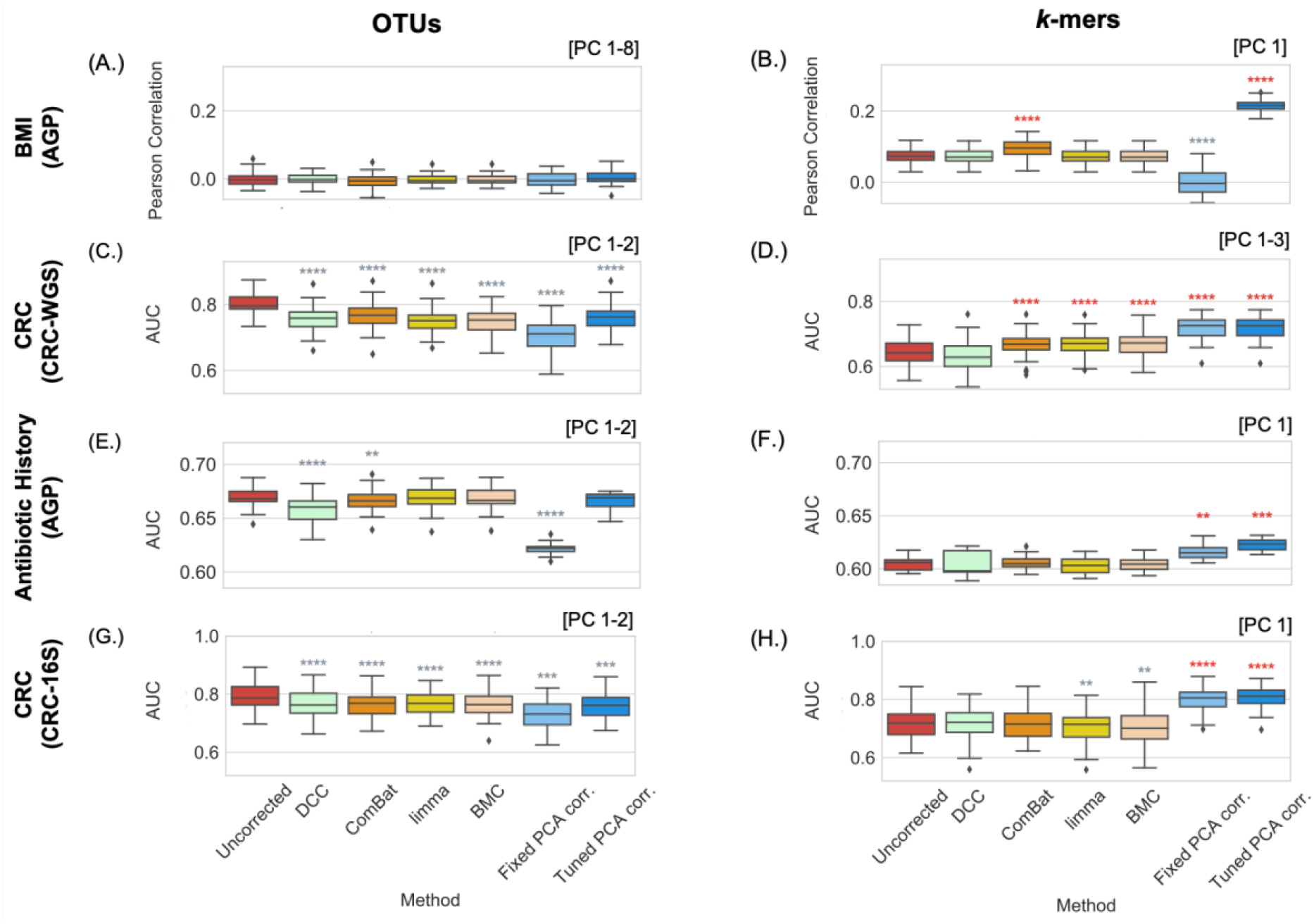
Impact of confounder correction approaches on phenotype prediction accuracy. The distribution of AUC or Pearson correlation in a cross-validated prediction model using either uncorrected data or data after applying one of the following covariate correction approaches: DCC, ComBat^51^, limma^50^, BMC^49^, and fixed-PCA correction with three PCs regressed out, and tuned-PCA correction where the number of PCs regressed out is a tuned hyperparameter. Distributions with a significantly higher mean compared to the uncorrected data are indicated with red asterisks, and distributions with a significantly lower mean compared to the uncorrected data are indicated with grey asterisks. The number of asterisks corresponds to the significance of a paired t-test between the uncorrected and corrected data (Methods). [ns,*,**,*** indicate *p*-values as follows: non-significant at 0.05 threshold, 10^−2^ < *p* < 0.05, 10^−3^ < *p* < 10^−2^, 10^−4^ < *p* < 10^−3^, *p* < 10^−4^, respectively, in a Wilcoxon rank sum test comparing each method to the baseline]. All methods’ baseline was Uncorrected except Fixed-PCA, which was compared to DCC as a baseline. Shown at top right is the number of PCs regressed out in the Tuned-PCA method

Among *k*-mers, supervised approaches significantly improved prediction accuracy of BMI in AGP and CRC status in CRC-WGS, but in the case of antibiotic history in AGP and CRC in CRC-16S, they had no effect or significantly reduced accuracy compared to no correction (**Figure 4**). By contrast, application of fixed and tuned-PCA correction to *k*-mers showed the greatest improvement in phenotype prediction for most datasets and phenotypes (**Figure 4**).

Although *k*-mers showed lower overall phenotype prediction accuracy compared to OTUs (**Figure 4**), PCA correction was able to make *k*-mers into a feasible predictive input feature that has comparable prediction accuracy to OTUs. Thus, in scenarios where technical covariates account for more microbiome variation than the phenotype of interest does, as observed with *k-*mers, using a background noise correction strategy can significantly improve phenotype prediction accuracy.

## Discussion

The ability to predict human phenotypes from metagenomic data is important to establish biomarkers of disease and the subsequent development of therapeutics. However, a major issue that impacts accurate prediction is the presence of confounders and systemic background noise both within^26^ and across studies^15,31^. In this paper, we investigated the ability of existing approaches to correct for sources of background noise in microbiome data and evaluated the potential utility of an unsupervised approach – PCA correction on CLR-transformed data.

PCA correction has been effective in correcting for unwanted variation in human genetic data and morphological data ^65–68^, but to date has not been applied to correct for such noise in microbiome data. Yet, we and others have found that top principal components in multiple datasets are correlated with numerous potential sources of unwanted noise such as host genetics^69^, ethnicity of the host^70^, and also abiotic factors like temperature^71^, suggesting that PCA correction may be an effective unsupervised correction approach. We found that regressing out the top PCs after applying a CLR transformation can address multiple issues simultaneously: first, this approach can reduce variance associated with technical and biological factors while improving the proportion of variance attributable to the phenotype of interest and resulting in improved ability to predict phenotypes (**Figure 4**). Additionally, the output from PCA correction can further be used for downstream analyses such as differential abundance analysis. We found that the extent of background noise differs from one dataset to another, and thus the ideal number of PCs to regress out with either a fixed or tuned PCA correction varies for each dataset. For example, the top five PCs in the AGP dataset are significantly correlated with technical variables, whereas the top nine PCs in the CRC-WGS are correlated with technical variables.

The application of CLR transformation can address many attributes of microbiome data that make it difficult to model including sparsity, non-normality (Figure S3) and dependency of features, which existing unsupervised approaches designed for non-microbiome data^43–45^ are ill-equipped to deal with. The application of CLR to microbiome data has been broadly recommended^53,72^ and is part of a suite of methods known as Compositional Data Analysis (CoDA)^73,74^ to address the dependency between features inherent to compositional data. However, the adoption of CLR in the microbiome field has not been uniform. Recently, McLaren at al.^15^ argued that CoDA methods can make microbiome data invariant to bias and suggests that it is underutilized within the field. Specifically, McLaren et al.^15^ found that that ratio-based analyses could remove intra-study bias, though did not address its effect on multiple datasets that are pooled together or large datasets with heterogeneous sampling procedures such as the AGP. Here, we provide the first systematic investigation into the effect of how CLR in combination with PCA can remove inter-study and intra-study bias. We hypothesized that applying CLR transformation will more readily reveal the covariates that introduce background noise across and within heterogeneous datasets because these contributors of bias (e.g. DNA extraction method, sequencing instrument, etc.) have a multiplicative effect on relative abundances ^15^. We found that indeed relationships between the microbiome and such variables is more apparent after CLR transformation (**Figure 2**), which is consistent with the multiplicative nature of bias expounded in McLaren et al.^15^ because the multiplicative bias becomes additive in log space, such that PCA is able to capture the bias in the top PCs as a shift in the centroid of samples plotted for a given dataset.

The supervised approaches^49–51^ are beneficial in that they directly remove confounding, potentially at the cost of phenotype prediction, while unsupervised approaches are beneficial in correcting for both measured and unmeasured factors of microbiome. Correcting for confounders naturally tends to result in removal of phenotype signal, as is the case in ComBat^51^ and fixed-PCA, which may be more effectively removing confounding effects at the cost of reduced proportion of variance explained by phenotype (**Figure 3**). Tuned-PCA protects the phenotype effect by removing up to the first PC that would significantly remove phenotype signal. However, caution must be taken when using tuned-PCA in the presence of strong confounding as it may not remove all confounding in order to protect the phenotype effect. In these scenarios, one should consider either a liberal correction of confounding by correcting for more PCs or subsampling the data such that cases and controls are matched for known confounders as is done in Vujkovic-Cvijin et al. ^26^.

In addition to optimizing the number of PCs for regression, we similarly identified the best Random Forest parameters and data transformations for the ComBat^51^, limma^50^, and BMC^49^ methods to fully evaluate the strengths of each approach (e.g. log transformed input was optimal for ComBat^51^). Note that we did not compare methods to percentile normalization^36^, which is a supervised approach recently proposed for robust biomarker discovery rather than phenotype prediction. Percentile normalization requires using all the control samples in one dataset to determine the percentiles for all the case samples in that same dataset, and so it is not immediately obvious how to apply this to a phenotype prediction model in which held out data is required. Future work adapting this method for phenotype prediction purposes may be an exciting area of methodological development.

We compared the prediction ability and impact of background noise correction methods on OTUs versus *k-*mers. *K-*mers are a convenient reference-free alternative to taxonomic features in that they produce non-sparse features that are also normally distributed at small values of *k*. Previous work has found that *k*-mers of sizes six and seven from microbiome data are predictive of phenotypes^75^. However, *k*-mers are limited because they are usually not directly interpretable biological features, although they can still be used as biomarkers of disease. This limitation may be a reason why OTUs outperform *k-*mers in phenotype prediction accuracy (**Figure 4**). It is crucial to note however, that *k*-mers may provide a better signature of technical artifacts like PCR bias^76,77^ and are also known to be protocol specific^78^. Thus, this may explain why for both 16S and WGS data, *k*-mers had higher correlations with technical variables compared to OTUs (**Figures 2, S2 and S5**). This aspect of *k-*mers offers justification for why PCA correction is particularly effective with *k*-mers.

As we hypothesized, in scenarios where the phenotype of interest is not directly correlated with other covariates, supervised approaches for batch correction and PCA correction yield better prediction than uncorrected data. This was true for *k-*mers where non-phenotypic variables were not as correlated with the phenotypes of interest compared to OTUs. This observation offers another justification for why PCA correction improved phenotype prediction in *k*-mers whereas in OTUs all correction methods reduced phenotype prediction. The difference in distribution of cases and controls between different studies is more drastic for some pooled datasets than others, reflecting a higher possibility of confounding (**Table S1**). We note that even though application of background noise correction approaches to OTUs does not always improve phenotype prediction accuracy, by applying correction, the true causal OTUs for the phenotype may be easier to identify with fewer false positives^31^.

Background noise correction is becoming increasingly important as the microbiome field matures and new datasets become available. One exciting future application of correction that we foresee is in microbiome wide association studies in which microbiome genomic polymorphisms are associated with human phenotypes^79,80^. Such a scenario may benefit from background noise correction since population structure may play a considerable confounding role^81^. As researchers consider the best approach for background noise correction for their specific research questions, they must weigh the tradeoffs between addressing confounding while also maintaining as much of the phenotype signal as possible. There is no single solution that will address all problems, but at minimum researchers should perform careful forensics to investigate the nature and pervasiveness of confounders in their data. In this manner, consistent and robust inferences can be made across multiple studies, moving us towards the goal of accurate phenotype prediction from microbiome data.

## Methods

### Datasets

Raw 16S fastq files were downloaded from the NCBI Sequence Read Archive (SRA) with study accessions PRJEB11419 for the American Gut Project, and PRJNA290926 (Baxter et al.^58^) and PRJEB6070 (Zeller et al.^82^) for CRC-16S. Fastq files for Zackular et al.^60^ from CRC-16S were obtained from http://mothur.org/MicrobiomeBiomarkerCRC/. The raw WGS fastq files for CRC-WGS were downloaded from SRA with study accessions PRJEB12449 (Vogtmann et al.^62^), PRJEB10878 (Yu et al.^63^), PRJEB7774 (Feng et al.^61^), PRJNA447983 (Thomas et al. Italian validation cohorts^1^), PRJEB6070 (Zeller et al.^82^), and PRJNA389927 (Hannigan et al.^64^). Processed OTU data for the AGP was obtained from Qiita study id 10317 (EBI submission <u>ERP012803</u>). OTU profiles from CRC-16S were obtained from the MicrobiomeHD database (Duvallet et al.^8^). Taxonomic profiles for CRC-WGS were obtained through the R package curatedMetagenomicData^83^ which used MetaPhlAn2^84^.

### Raw Sequence Processing

Features in metagenomic data can be defined in two broad ways, both high-dimensional: reference-based approaches and reference-free approaches. Reference-based approaches cluster sequenced reads based on a defined threshold and assign taxonomy by aligning reads to reference genomes. Reference-free approaches, sort reads into bins that are defined independently of known genomes, i.e. *k*-mers, short strings of length *k* that can be obtained directly from read sequences, which are increasingly popular in microbiome data analyses and have been used by several studies to do prediction ^85–88^. *K*-mers offer a powerful alternative approach to more commonly used Operational Taxonomic Units (OTUs) because they do not rely on a reference database of genomes and do not require identifying a set of parameters to determine OTUs ^89^.

To compute *k*-mer abundances, raw sequences were input into the *k*-mer counting algorithm Jellyfish 2.3.0^90^ with default parameters except for a hash of 10 million elements and canonical *k*-mers with size of 5,6,7 or 8. Prior work has shown that *k*-mer sizes of 6 and 7 are predictive of phenotype^56^. The resulting *k-*mer abundance table is then converted to a composition such that each sample sums to 1 to account for different reads depths across samples. Taxonomic profiles were similarly converted to compositions.

### Centered log ratio transformation

The centered log ratio (CLR) transformation is a compositional data transformation that takes the log ratio of between observed frequencies and their geometric means. This is done within each sample where relative frequencies of different taxa are measured and sum to 1. This can be written in mathematical form as:

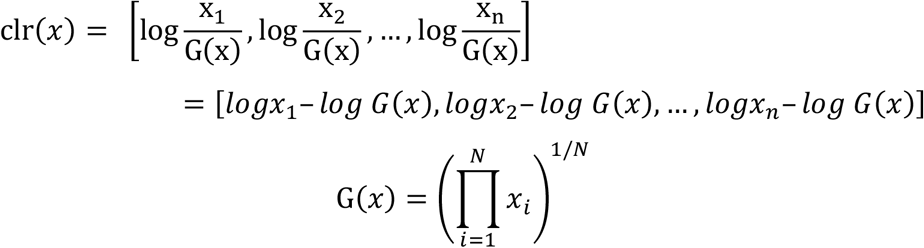

Here, ***x*** is a vector representing the abundance of microbiome features in a single sample, and *G*(***x***) represents the geometric mean. Prior to applying PCA, the CLR-transformed values are standardized such that all samples have a mean of 0 and variance of 1. The Gaussian-like distribution of CLR-transformed microbiome compositional data is shown in **Figure S3**.

### Background noise correction methods

The existing supervised approaches for background noise correction compared in this study include batch mean centering (BMC)^49^, ComBat^51^, and limma^50^ applied to relative abundance data. ComBat^51^ assumes data is cleaned and normalized prior to batch effect removal. It’s common to add a pseudocount of 1 to 0 observations so that one can apply a log transform in the normalization prior to ComBat^51^ (as described in Gibbons et al.^31^). We followed this same procedure with both OTU and *k*-mer, and applied ComBat^51^ to the log of relative abundance data. For limma, batch mean centering (BMC), and Direct Covariate Correction (DCC) we used the relative abundance. The CLR transformation and PCA-Correction used the relative abundance of *k*-mers and taxonomic features. The equation used to regress out confounding covariates in DCC is as follows:

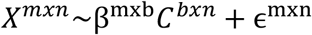

Where the original feature matrix *X* with *m* features and *n* samples is the outcome of a linear model with covariate associated coefficient matrix *β*, dummy matrix *C* with each row representing one of the *b* possible values of the confounding covariate, and *ϵ*, the residual matrix. The residual matrix *ϵ* is the covariate-corrected feature matrix.

In PCA correction, top PCs computed from the CLR transformed *k*-mer or OTU relative abundance tables are regressed out. The CLR transformation cancels out the multiplicative bias^15^ within each study by taking a ratio of features to the geometric mean of features that are all impacted by the same study-specific multiplicative bias. The transformation accentuates the difference in bias across studies by smoothing out the intra-study bias, thereby allowing PC regression to account for the confounding across studies. In the fixed-PCA correction, a set number of PCs are regressed out from the microbiome data. In the main figures we show results after regressing up to three PCs. We show results regressing up to 9 PCs in the supplement. Alternatively, the tuned-PCA correction uses a train-validation-test approach to tune two hyperparameters: the optimal number of PCs to regress out *p*, and, when using *k*-mers, the optimal *k*. The same portion of data used for validation in the Random Forest tuning is used for tuning the PCA correction hyperparameters, where the tuned Random Forest hyperparameters are fixed before tuning *p* and *k*. To determine the number of PCs that optimize phenotype prediction, PCs 1 through *p* were regressed out of the input data with *p* ranging from 1 to 20. The *p* that produces the highest AUC or Pearson correlation in phenotype prediction (method of prediction model described below) in validation was selected. The same procedure is done with *k* where values between 5 and 8 are tested (only *k*-mer sizes 6 and 7 were tested for CRC-WGS) The reported performance is based on the remaining 20% set aside for testing.

### Variance components analysis

We performed variance components analysis before and after correction to show how batch correction impacts biological signal of interest and technical signal. Specifically, a linear mixed model was fit to the data where the outcome was abundance assigned to a taxa or *k*-mer, and the variables of the model were various biological and technical variables obtained from a study’s metadata. The outcome was represented by a vector (with dimensions *t* features x 1), and the biological and technical covariates were represented by a design matrix (with dimension of number of covariates *c* x number of samples *n*). Continuous variables, e.g. BMI, and categorical variables with two or less levels (e.g. sex) were modeled as fixed effects while categorical variable with more than two levels (e.g. ethnicity) were modelled as random effects. This LMM model was computed with the R package variancePartition^91^. For example, in the AGP, the following LMM was implemented:

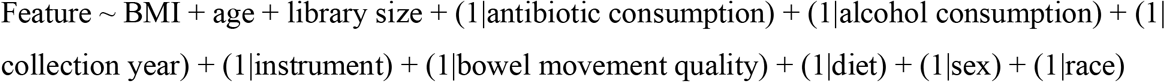

For AGP, sequencing instrument, collection year, race, alcohol consumption, diet, bowel movement and antibiotic consumption were modeled as random effects while library size, age, and BMI were modeled as fixed effects. For CRC-WGS, colorectal cancer status, dataset label, library layout, sequencing instrument, sequencing center, and DNA extraction kit, and sex were modeled as random effects and library size was modeled as a fixed effect. For 16S, dataset label, sequencing method, race, CRC status, and sex were modeled as random effects while library size and age were modeled as fixed effects.

### Phenotype prediction

In CRC-16S and CRC-WGS, we predicted whether a sample comes from a host with colorectal cancer or a healthy host. For the American Gut Project, we predicted whether a sample comes from a host who took antibiotics in the previous year or a host who has not taken antibiotics in the previous year. We also use the American Gut Project to predict body mass index (BMI).

We performed prediction of binary traits using Random Forest implemented in Scikit-learn^92^, which has been previously employed successfully for predicting binary outcomes from microbiome data^32,34,93,94^. We tuned four hyper-parameters of the Random Forest model in a grid search using a train-validation-test strategy with 56% of samples in the meta-cohort used for training, 24% for validation of model hyper-parameters, and 20% reserved for testing. These four hyperparameters included the number of estimator trees (100, 1000, or 1500), criterion (entropy or gini), minimum samples per leaf (1, 5, or 10), and features (0.1, 0.3, 0.5). Minimum samples per split was set at 2 and max depth was set at 5, and default parameters otherwise. This was performed in five-fold cross validation repeated ten-times to obtain confidence intervals on the area under the ROC curve (AUC), our metric of prediction accuracy. A similar train-validation-test strategy was used for the linear regression model to select coefficients of the model where accuracy was measured using Pearson correlation of the true BMI to the predicted BMI. The difference in the distribution of prediction accuracy for both prediction tasks was quantified statistically using a paired Student’s t-test.

### Code Availability

An implementation of PCA correction and phenotype classification is available at https://github.com/garudlab/Microbiome_PCA_correction.

## Acknowledgements

We thank members of the Halperin Lab and Garud Lab, as well as Michael R. McLaren for helpful discussions.

## Author Contributions

Conceived and designed the experiments: L.B., B.B., S.S., E.H., N.R.G. Performed the experiments: L.B., B.B. Analyzed the data: L.B., B.B. Coded the pipeline: L.B. Contributed analysis tools: E.H., N.R.G. Wrote the paper: L.B., B.B., S.S., E.H., N.R.G. All authors have read and approved the manuscript.

