## Supplementary Figures and Tables for "Correcting for Background Noise Improves Phenotype Prediction from Human Gut Microbiome Data"

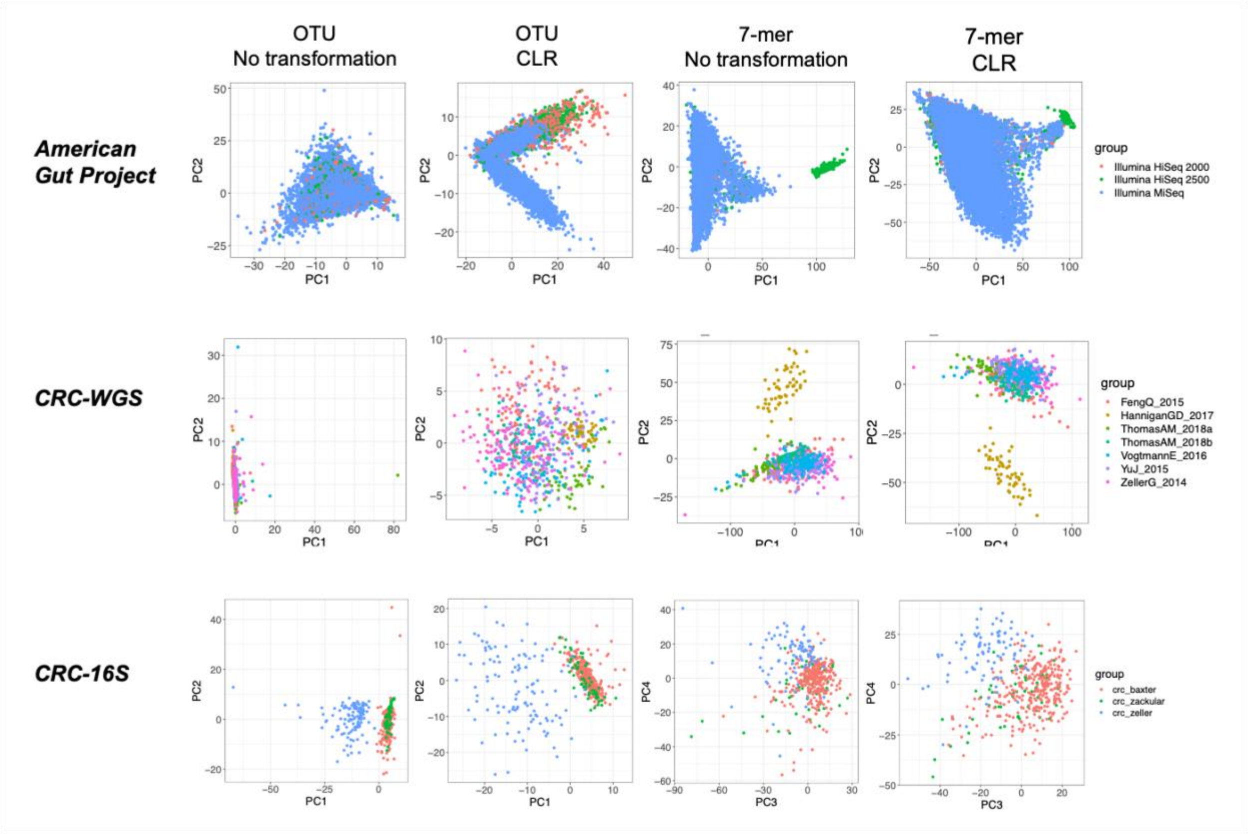


**Figure S1. First two principal components from microbiome dataset studied.** PCA was applied to OTU and 7-mer data from the AGP, CRC-WGS merged dataset, and CRC-16S merge datasets. Samples were plotted along the first 2 PCs with colors indicating dataset membership.

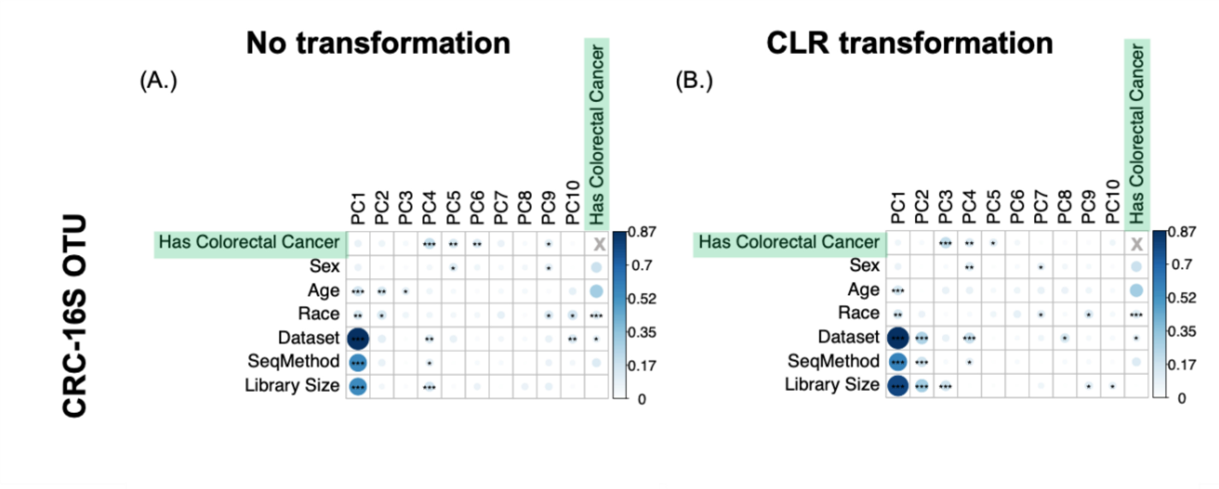


**Figure S2. Top principal components from the CRC-16S dataset correlate with technical and biological covariates.** The first 10 PCs in the CRC-16S OTU joined dataset are correlated with variables measured in each of the studies, including colorectal cancer status (CRC), sex, age, race, dataset label, sequencing method, and library size. The size and color of the circles in each cell indicate the magnitude of correlation while black asterisks indicate the significance of the Pearson correlation of the PCs with each of the variables. The color bar at right of each plot represents the range of correlations observed across all datasets. [*,**,*** indicate *p*-values as follows: $10^{-2} < p < 0.05$, $10^{-3} < p < 10^{-2}$, $10^{-4} < p < 10^{-3}$, $p < 10^{-4}$].

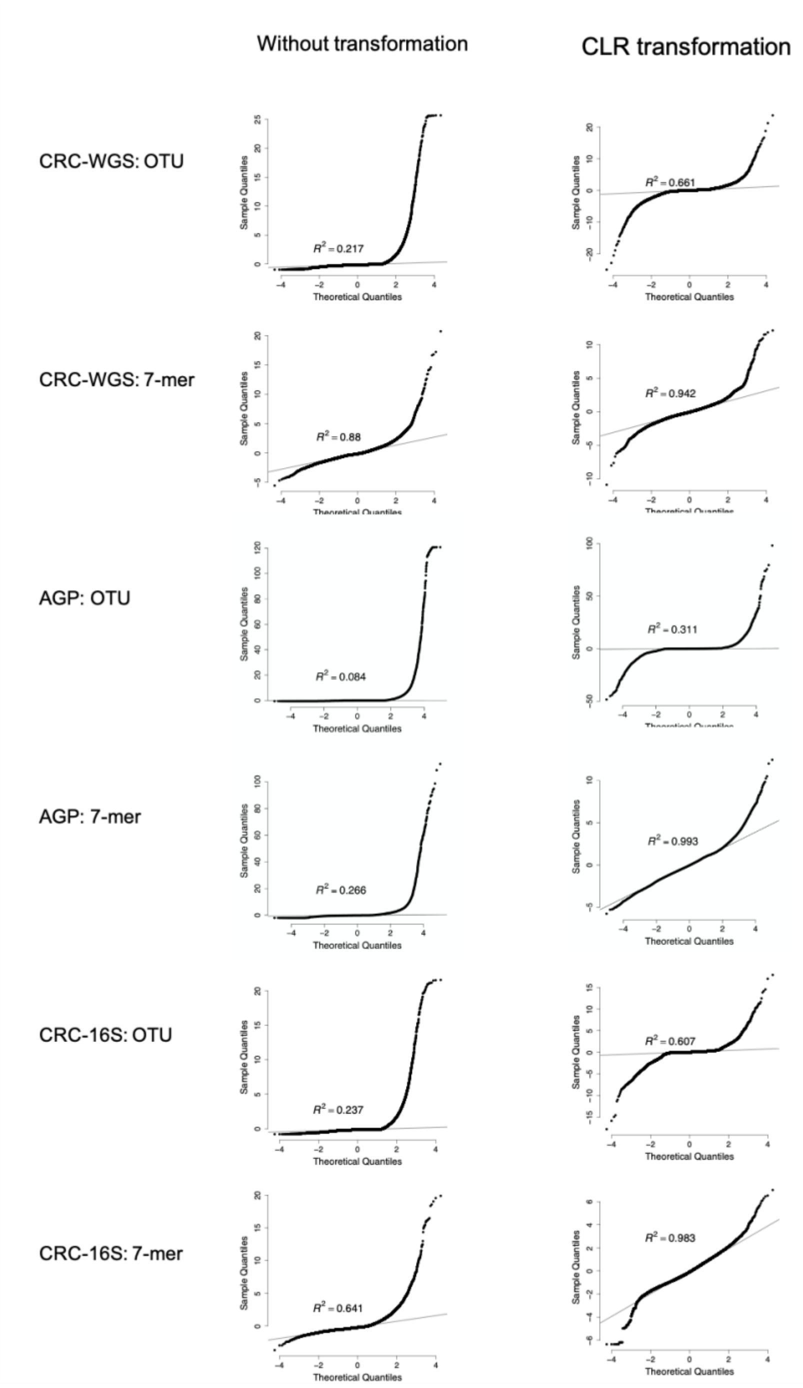


**Figure S3. Quantile-Quantiles plot for AGP, CRC-WGS, and CRC-16S before and after the CLR-transformation.** The quantiles of 100 randomly-selected OTU features or *k*-mers, that were converted to z-scores, ranked against the expected quantiles from a normal distribution of mean 0 and variance 1. The R-squared values are reported in the annotated text.

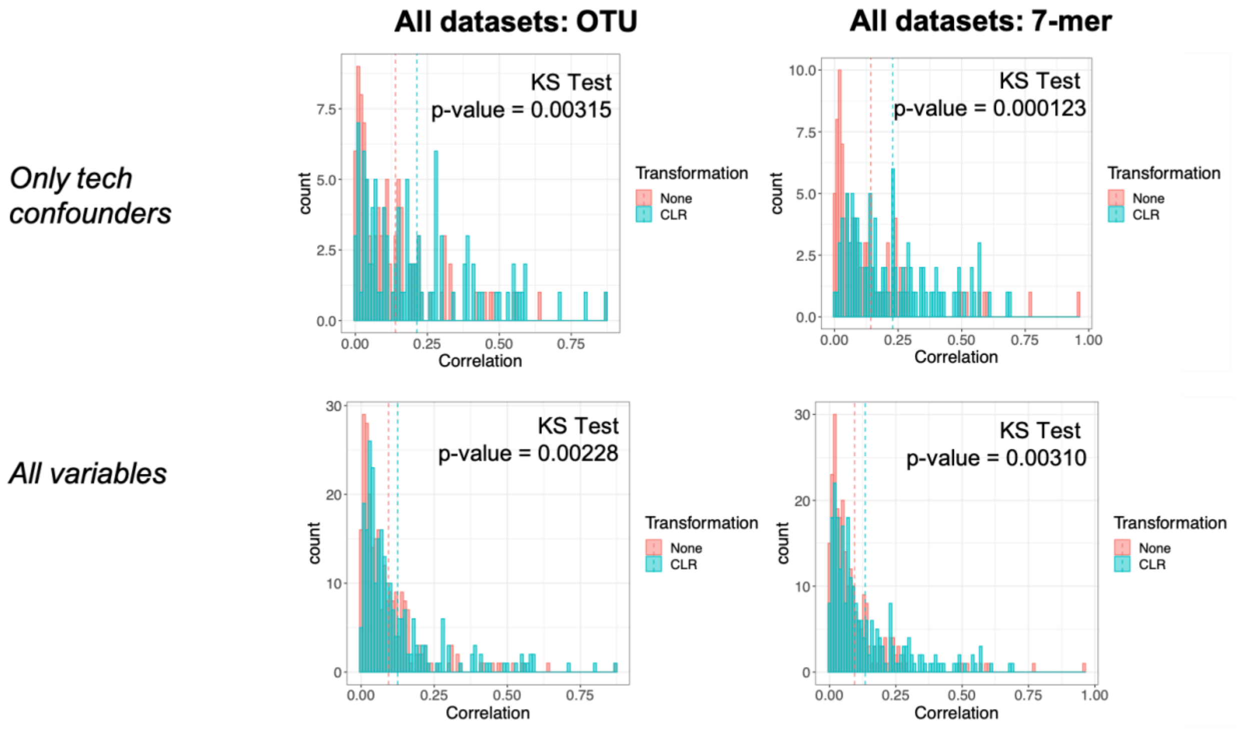


**Figure S4. Histogram of correlation between top 10 PCs and technical confounders or all measured variables.** Shown are the Kolmogorov-Smirnov test p-values for the test of the null hypothesis that the correlations of the top 10 PCS in the non-transformed data is from the same distribution as the CLR-transformed data. In both OTUs and 7-mers, the CLR-transformed data have a significantly higher distribution of correlations with top PCs compared to the non-transformed data.

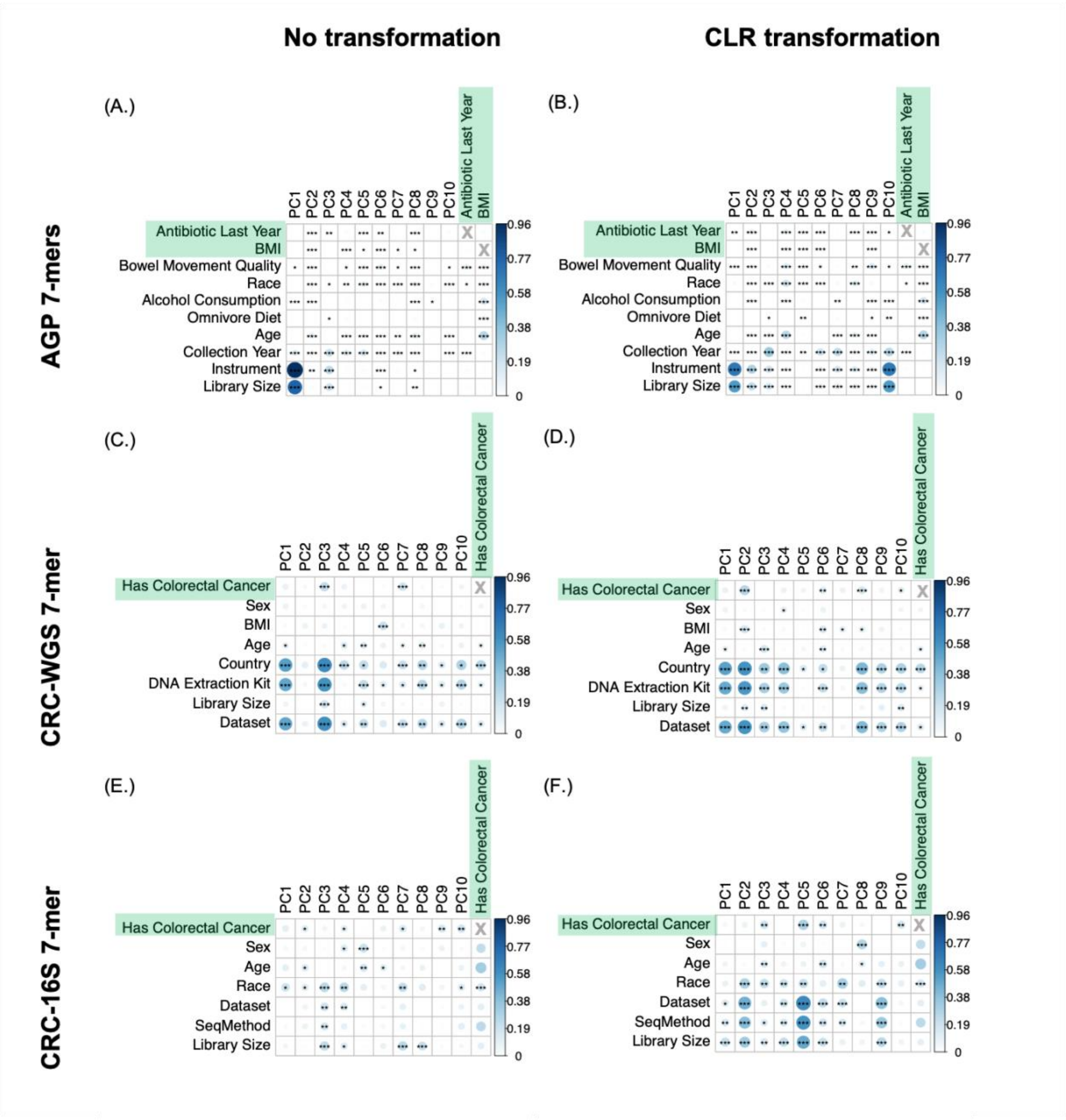


**Figure S5. Top principal components from 7-mers correlate with technical and biological covariates.** The first 10 PCs before (a, b, and c) and after (b, d, and f) the CLR-transformation are correlated with variables measured in each of the studies, including dataset label, library size, DNA extraction kit used, country of origin, age, body mass index (BMI), sex, and colorectal cancer status (CRC). The size and color of the circles in each cell indicate the magnitude of correlation while black asterisks indicate the significance of the Pearson correlation of the PCs with each of the variables. The color bar at right of each plot represents the range of correlations observed across all datasets. [*,**,*** indicate $p$-values as follows: $10^{-2} < p < 0.05$, $10^{-3} < p < 10^{-2}$, $10^{-4} < p < 10^{-3}$, $p < 10^{-4}$].

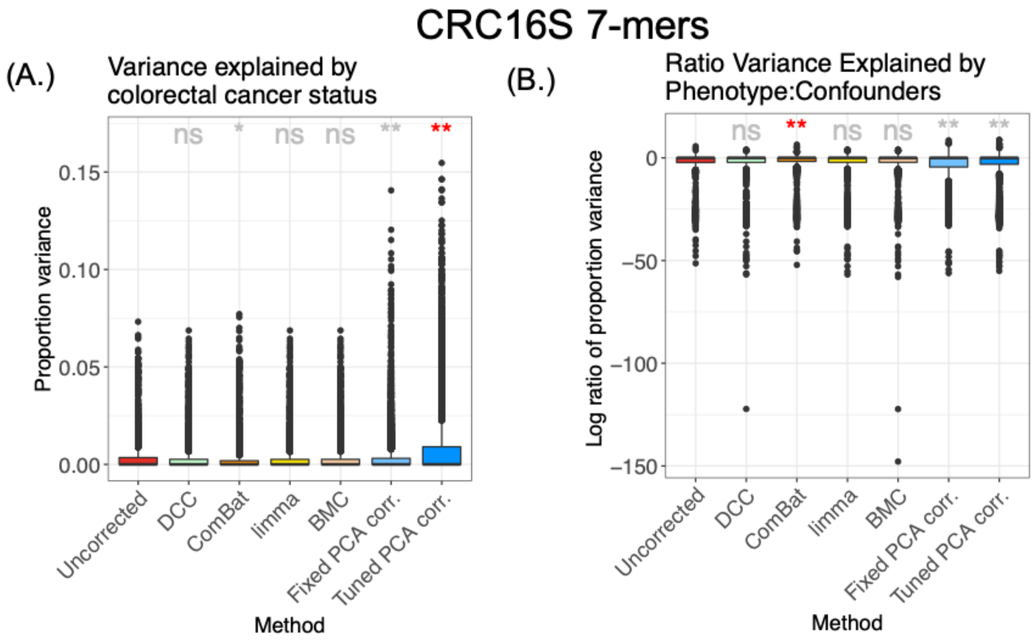


**Figure S6. Proportion of variance explained by phenotype vs. confounders.** We plot the distribution of the proportion of variance explained by (a) colorectal cancer status in the CRC-16S joined datasets. We then plot the log2 ratio of that same proportion by the proportion explain by confounders including dataset label, sequencing method, race, sex, and library size in the CRC-16S joined dataset. For all plots, each data point represents one feature in the 7-mer matrix table. Distributions with a significantly higher mean compared to the uncorrected data are indicated with red asterisks, and distributions with a significantly lower mean compared to the uncorrected data are indicated with grey asterisks. [ns,*,**,*** indicate $p$-values as follows: non-significant at 0.05 threshold, $10^{-2} < p < 0.05$, $10^{-3} < p < 10^{-2}$, $10^{-4} < p < 10^{-3}$, $p < 10^{-4}$, respectively, in a Wilcoxon rank sum test comparing each method to the baseline].All methods' baseline is Uncorrected except Fixed PCA, which is compared to DCC as a baseline.

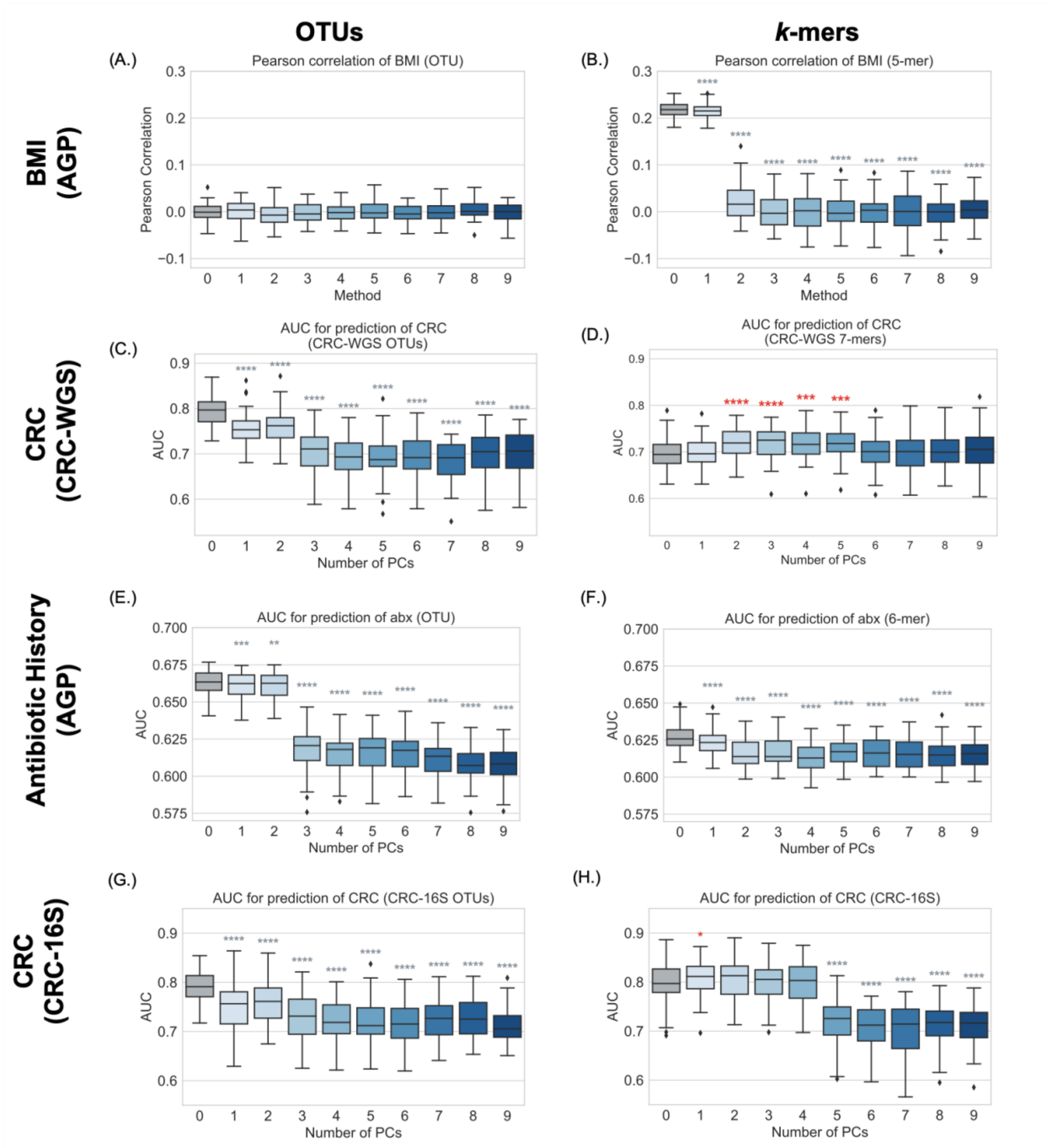


**Figure S7**. **Regressing out top PCs from CRC-WGS 7-mer data improves prediction of colorectal cancer.** The x-axis indicates the number of PCs regressed from the 7-mer matrix, where 0 indicates no correction and 1 to 10 represent 1 to 10 PCs regressed out from the 7-mer matrix. The y-axis represents the AUC of predicted CRC-status. Distributions with a significantly higher mean compared to the uncorrected data are indicated with red asterisks, and distributions with a significantly lower mean compared to the uncorrected data are indicated with grey asterisks. [ns,*,**,*** indicate *p*-values as follows: non-significant at 0.05 threshold, $10^{-2} < p < 0.05$, $10^{-3} < p < 10^{-2}$, $10^{-4} < p < 10^{-3}$, $p < 10^{-4}$, respectively, in a Wilcoxon rank sum test comparing each method to the baseline].

| Phenotype | Joined dataset | Batch/dataset | Number of cases | Number of controls |
|---|---|---|---|---|
| Antibiotic history | AGP[1] | Illumina Hi Seq 2000 | **186** | **301** |
| | | Illumina Hi Seq 2500 | **312** | **607** |
| | | Illumina MiSeq | **5,083** | **10,457** |
| Colorectal Cancer | CRC-16S | Baxter et al.[2] | **120** | **172** |
| | | Zackular et al.[3] | 30 | 30 |
| | | Zeller et al.[4] | **41** | **75** |
| Colorectal Cancer | CRC-WGS | Feng et al. 2015[5] | **46** | **61** |
| | | Hannigan et al. 2017[6] | 27 | 28 |
| | | Thomas et al. 2018a[7] | 29 | 24 |
| | | Thomas et al. 2018b[7] | 32 | 28 |
| | | Vogtmann et al. 2016 [8] | 52 | 52 |
| | | Yu et al. 2015[9] | **75** | **53** |
| | | Zeller et al. 2014[4] | **91** | **66** |

**Table S1. Number of cases and controls for binary phenotypes per study included in joined dataset.** For each binary phenotype of interest in the AGP, CRC-WGS, and CRC-16S, we show the number of cases and controls for each respective subset of the data based on sequencing instrument (AGP) or original source study (CRC-WGS and CRC-16S).
